# Neonatal Platelets Differentiate Monocytes to a Myeloid Derived Suppressor Cell Phenotype

**DOI:** 10.1101/2025.11.21.689809

**Authors:** Preeti Maurya, Daniel O’Reilly, Zachary T. Hilt, Alison C. Livada, Kathleen E. McGrath, Chen Li, Sara K. Ture, Michael W. Malloy, Ei Thanda Tun, James Palis, Craig N. Morrell

## Abstract

Adult platelets are relatively enriched in immune related molecules compared to neonatal platelets, but neonatal platelets express some growth factors and enzymes at comparatively higher levels. This makes a prediction of platelet-immune cell interaction outcomes in neonates a challenge, as they are likely dependent on the cell type and tissue environment at the time of injury or infection. Our past studies revealed that the transfusion of adult but not neonatal platelets into thrombocytopenic neonatal mice led to an acute increase in monocyte trafficking. We have now found that neonatal, but not adult platelets, induce monocytes to a Myeloid Derived Suppressor Cell (MDSC) phenotype, typified by increased PD-L1, that limits T-cell activation *in vitro* and *in vivo*. Monocytes prior incubated with neonatal, but not adult platelets, or platelet releasates, limited T-cell activation *in vitro*. Using an *in vivo* asthma-like model we also found that the treatment of mice with monocytes prior incubated with neonatal platelet releasates limited T-cell activation in a asthma-like model. Platelet-driven effects were dependent on neonatal platelets producing more PGE_2_ that signaled through monocyte EP4. These studies indicate that neonatal platelets have immune limiting roles in the post-natal period by indirectly limiting T-cell responses, perhaps contributing to the adverse outcomes of platelet transfusions to neonates.

## Introduction

Neonates and adults have different pathogen responses^1,2^. Neonates have an immune system adapted to address the unique challenges during a time of continued tissue development and microflora colonization. Their responses depend on the type of pathogen, the stage of pre/post-natal development, and a multitude of environmental and maternal factors^3–5^. To avoid excessive or mis-directed neonatal immune activation, while also providing protection, the maternal immune system participates in pathogen responses in part through the transplacental transfer of maternal IgG and the transfer of IgA in breast milk^2^. Myeloid Derived Suppressor Cells (MDSCs) are a set of myeloid lineage cells more numerous in neonates compared to adults with an immune limiting function, particularly in limiting T-cell activation^6^. MDSC numbers decline with post-natal age and are nearly absent in healthy adults^6,7^. MDSCs are divided into two groups depending on their cell lineage origin; monocytic-myeloid derived suppressor cells (M-MDSCs) and polymorphonuclear-myeloid derived suppressor cells (PMN-MDSCs)^8^.

MDSCs are typically seen as having adverse effects in the adult by limiting infection and tumor responses, creating an immune-tolerant microenvironment that may prolong or worsen disease outcomes^9^. The expansion of MDSCs in adults is regulated by STAT family transcription factors (STAT3, STAT5 and STAT6) with the help of Arginase-1^10,11^, as well as by hypoxia inducible factors (Hif1α)^12^. In some conditions platelets can mediate immune suppressive cell differentiation via secreted factors such as platelet activating factor (PAF) and TGF-β ^13,14^. Fatty acid transporter protein 2 (FATP2) regulates MDSCs differentiation upstream of arachidonic acid and PGE_2_ synthesis, that ultimately increases MDSCs ^15^.

Platelets are not only the cell mediators of thrombosis, but platelets also directly and indirectly regulate all phases of immune responses^16–18^. A decline in platelets (thrombocytopenia) can result in immune dysfunction, while platelet activation results in immune activation^19,20^. Platelets are therefore rheostats of vascular inflammation and central regulators of immune responses. Adult and neonatal platelets differentially express many immune related molecules, including neonatal platelets having less P-selectin, β2M, PDGFBB, SDF-1 and complement proteins^20–24^, but there has been little functional analysis of how this affects immune development and responses. Thrombocytopenia is a common complication of prematurity and the transfusion of adult platelets to neonates is associated with multiple adverse outcomes, including increased risk of mortality^25,26^. Pre-term neonates treated at a higher transfusion threshold (50,000 platelets/µL) had significantly higher rates of death or major bleeding complications compared to neonates given transfusions at a lower threshold (25,000 platelets/µL)^27^. Our prior studies using a mouse model of acute neonatal platelet transfusion demonstrated that the adverse effects of adult platelet transfusions may in part be due to the impacts of the immune composition of adult and neonatal platelets on monocyte trafficking^21^. Because the cell, tissue, and developmental environments of adult and neonates are so different and continue to change, the immune function of adult and neonatal platelets must develop to meet challenges in an age-appropriate manner.

In the present study, we discovered that neonatal and adult platelets induce different patterns of monocyte differentiation. Not only do neonatal platelets increase the expression of some inflammatory markers such as Ly6C, but neonatal platelet-derived products induce monocyte differentiation to immune suppressive M-MDSCs that limit T-cell activation. Neonatal platelet induced M-MDSCs may in part be due to a higher production of PGE_2_, leading to a limiting of adaptive immune functions.

## Methods

### Animal studies

Experimental protocols were approved by the URMC Institutional Animal Care and Use Committee. All mice were on a C57BL/6J background. Both male and female mice were used for all mouse experiments and platelets from post-natal day 1 (P1) to P3 and 8-12 weeks old mice were used. Six week old C57BL/6J mice were used for myeloid derived suppressor cells (MDSCs) transfer experiments. P1-P3 pups were euthanized followed by blood collection into heparinized Tyrode’s buffer to isolate neonatal platelets. For adult platelets, mice were bled via retro-orbital collection into heparinized Tyrode buffer followed by platelets isolation as prior described^20,21^. To prepare releasate, platelets were resuspended in buffer and treated with thrombin (2U/ml) for 10 minutes. Hirudin (2U/ml) was then added to neutralize thrombin. Platelets were centrifuged at 100G for 10 minutes and the supernatant collected as the platelet releasate. 1x10^6^ bone marrow monocytes were seeded in each well and platelet releasate added as 1 monocytes:resleasate from 20 platelets, and cells harvested after 72 hr.

For allergy induced airway inflammation mouse models, 6 week old mice were used. Adult and neonatal platelet releasates were co-cultured with monocytes for 3d and 3x10^6^ cells intravenously infused (retro-orbital) into mice on d0 and d6, followed by alum-OVA administration (intra-peritoneal) on d1 and d7. To induce an allergic lung response, mice were administered ovalbumin (OVA) oropharyngeal on d14, 15, and 16. OVA was dissolved in normal saline and 5 mg/kg administered.

### Mass Spectrometry

Neonatal and adult platelets were collected into heparinized Tyrodes. After platelet isolation by centrifugation, platelets underwent a negative selection procedure to ensure purity using biotin labelled anti-CD45, CD220, Ter119 followed by addition of Streptavidin Dynabeads. After incubation at room temperature the tubes are placed into a magnetic stand and the supernatant poured off. Platelets were lysed using RIPA lysis buffer and the samples were collected for mass spectrophotometry analysis. Mass spectrometry preparation was performed by the mass spectrometry core facility in the University of Rochester as per local protocol using S-Trap sample processing kits (ProtiFI, USA) with 500ng of protein loaded for each run. Samples were analysed using an Orbitrap Astral Mass Spectrometer (Thermo Fisher Scientific, USA) with data independent acquisition (DIA) applied to identified peptide peaks using the Astral analysis platform.

### Cell culture

Bone marrow monocytes were isolated from 8-12 week old mice using a Stem Cell EasySep^TM^ mouse monocytes isolation kit (catalog no. 19861) per manufacturer instructions. After isolation, platelets were activated with thrombin (2 U/ml) for 10 minutes followed by neutralization of thrombin by adding equal concentration of Hirudin (2 U/ml). Platelets were then centrifuged at 2600 rpm for 10 minutes and the supernatant collected (called platelet releasate). For our experiments, we treated the monocytes with platelet releasates from 20 platelets for every one monocyte (20:1 ratio) for 72 hr. Cell culture media was supplemented with G-CSF (2 nM) and GM-CSF (2 nM) to facilitate monocyte survival.

EP2 blocker (PF-04418948, Cayman Chemical) and EP4 blocker (ONO-AE3-208 Cayman Chemical) were used to block Prostaglandin E_2_ (PGE_2_) signaling. 18 µM of blockers were incubated with monocytes for 10 minutes before the platelet releasate treatment. To check the effect of PGE_2_ on monocyte phenotype, recombinant PGE_2_ (Item number 18220, Cayman Chemical) was used.

### T-cell and MDSCs Co-culture Assays

T-cells were isolated using a Stem Cell Technology total T-cell isolation kit (cat no. 19851) as per manufacturer instructions. To measure the proliferation of T-cells, freshly isolated T-cells were incubated with CFSE (5 µM) dye for 20 minutes. After 20 minutes, T-cells were washed twice with PBS. For T-cell activation, cells were incubated with anti-CD3/CD28 beads.

### Flow cytometry

Monocytes were washed and incubated with anti-mouse CD11b, Ly6C, PD-L1, DR5 and/or Ly6G antibodies as well as 7AAD as a live/dead marker for 30 minutes. Cells were washed and fixed with 1% formalin and data was acquired on LSR II flow cytometry machine. For T-cell activation and proliferation study, the co-cultured MDSCs and T-cells were collected on 5th day and spun down in 96 well V bottom plates and washed twice with PBS. After washing, cells were stained with anti-CD3, CD4, CD8, CD69, CD25 and/or 7AAD for 30 minutes. Cells were washed twice with PBS and fixed with 1% formalin and data was acquired on LSR II flow cytometry machine. All the data were analyzed using FlowJo software (version 10.7.1).

For the lung cell staining, lungs were mechanically minced using a syringe plunger and filtered through a 100 µm cell strainer. For RBCs lysis, ACK lysis buffer was used for 10 minutes followed by washing twice with PBS. Cells were stained with anti-CD45, CD3, CD4, CD8, CD25, CD69, Ly6G antibodies and/or 7AAD for 30 minutes at room temperature. Cells were washed twice with PBS and fixed with 1% formalin and data acquired on LSR II flow cytometry.

### Quantitative real-time PCR (qPCR)

Bone marrow monocytes isolated by negative selection were treated with neonatal or adult mouse platelets releasate for 72 hr. Monocytes were washed with PBS and lysed in RLT lysis buffer. RNA isolation was performed by the RNeasy mini kit (Qiagen). NanoDrop^TM^ 2000 (Thermo Fisher Scientific) was used to measure the concentration of mRNA and cDNA conversion was performed with a high-capacity RNA to cDNA kit (Applied Biosystem). Real time PCR was executed using TaqMan Gene Expression Master Mix Protocol and a CFX Connect Real-Time PCR Detection System (Bio-rad Thermocycler). TaqMan gene expression primer (Thermo Fisher Scientific, *Nos2* (Mm00440502_m1), *Arginase1* (Mm00475988_m1), *Cd274* (Mm03048248_m1), *Cxcl1* (Mm04207460_m1) and *Gapdh* (Mm99999915_g1) was used for the cDNA amplification in this study. The qPCR steps consist of 3 steps: Initiation (95^0^C for 10 minutes), denaturation (95^0^C for 15 sec) and annealing/extension (60^0^C for 1 minute). Amplification consists of denaturation and annealing steps and in this study, target cDNA was amplified for 40 cycles. Data analysis was done using calculation for fold change 2^(-ΔΔCT)^ with the housekeeping gene *Gapdh*. Samples were normalized with control and fold change values were reported.

### RNA-seq

For RNA-seq analysis, total mRNA isolation was performed using RNeasy mini kits (Qiagen kit). To digest genomic DNA, mRNA was treated with DNase and processed for RNA-seq with TruSeq Stranded mRNA library preparation. RNA-seq analysis was performed by University of Rochester Genomic Research Core and R-4.0.2 was used for data normalization and differential expression analysis.

### PGE_2_ ELISA

PGE_2_ was measured in adult and neonatal platelet releasates using a PGE_2_ ELISA kit (Item number: 514010, Cayman Chemical, USA) as per the manufacturer protocol. In brief, equal numbers of platelets isolated from adult and neonatal blood and whole platelets or platelet releasate prepared. Myassay software was used for data analysis.

### Immunoblot

Equal numbers of neonatal and adult platelets (5x10^6^) were lysed using RIPA lysis buffer followed by heat inactivation. The protein samples in the Laemmli buffer were loaded into Mini-PROTEAN® TGX Gels (BioRad). The loaded gels were run at 100V in 1X Tris/Glycine/SDS buffer. Transfer from the SDS-PAGE gel to nitrocellulose membrane (BioRad) was conducted at 80V for 2 hr with ice packs. Transferred blots were blocked in 5% BSA (Sigma Aldrich) in tris-buffered saline (TBS, Fisher Scientific) with 0.1% Tween-20 (TBS-T) for 1 hr, shaking at room temperature. Primary antibodies (PGES-1 and GAPDH) were diluted 1:1000 in 2.5% BSA and incubated overnight at 4^°^C. Anti-rabbit HRP was used as a secondary antibody (GE Healthcare) and diluted 1:2000 in 2.5% blocking buffer for 2 hours at room temperature with gentle agitation. Incubated membranes were developed using Supersignal West Pico (Thermo Fisher) using BioRad ChemiDoc MP chemiluminescence setting.

### Immunohistochemistry and Immunofluorescence

OVA administered mouse lungs were collected and placed in tissue fixative (60% MeOH, 10% acetic acid, 30% distilled water) then paraffin embedded and sectioned. For immunostaining, slides were deparaffinized and rehydrated and incubated in antigen retrieval solution for 20 minutes in a pressure cooker. After blocking, tissue sections were stained with primary antibodies GATA3 (1:250), T-bet1 (1:500) and incubated overnight at 4°C. After incubation, slides were washed in PBS 4 times and stained with ImmPRESS horseradish peroxidase anti-rabbit IgG reagent (Vector Laboratories) and were incubated for 30 minutes at room temperature. Slides were washed and stained with 3,3’ diaminobenzidine peroxidase substrate (Vector Laboratories) for 1-3 minutes. Slides were washed in distilled H_2_O for 5 minutes, counterstained with hematoxylin and dehydration performed. For imaging, an Olympus BX51 upright microscope was used. Analysis was performed using ImageJ and GraphPad Prism was used for statistical analysis.

For immunofluorescence, freshly isolated platelets were plated on poly-L-lysin coated glass chamber slides. Platelets were washed twice with Tyrode and fixed using methanol and acetone under chilled conditions for 5 minutes in each solution. The platelets were again thoroughly washed with PBS, and permeabilized with 0.1% Triton x-100 (in PBS) for 15 minutes followed by washing with PBS. 1% BSA in PBS was used for blocking at room temperature for 30 minutes. Platelets were stained with anti-Prostaglandin E synthase (PGES-1) antibodies (1:250; Abcam, USA #ab62050) at 4°C overnight followed by 4 times washing with PBS and staining with anti-rabbit Alexa Fluor 488-conjugated antibody for 2h at RT. After washing with PBS, nuclei were stained with DAPI (1μg/ml) and again washed twice with PBS. Platelets stained with anti-rabbit Alexa Fluor 488 were used as negative Control. Images were acquired for PGES-1 and DAPI staining using Olympus FV1000 confocal laser scanning microscope using a 60x oil-immersion objective lens with 8x electronic zoom and image resolution size was 1024x1024^24^. Image J software was used for the quantification of migratory cell number.

### Statistical analysis

Flow cytometry analysis was performed using the FlowJo, version 10.7.1, software and GraphPad Prism Software (La Jolla, CA, version 8.0) used for preparing graphs and all statistical analysis. A Shapiro-Wilk test was performed to check the normality of data when N was ≥6. For data that passed the normality test, and had >2 groups, a 1-way ANOVA was used with a Bonferroni multiple comparison correction. If the data sets did not pass the normality test, the Kruskal-Wallis test with Dunn multiple comparison correction was performed. For 2 groups comparison, Student *t* test was used. *P* values ≤0.05 were considered as statistically significant. All the data points represent biological replicates. Platelet proteomes were analyzed using R statistical software (Version 4.2.2, The R foundation for statistical computing, Vienna, Austria). Proteomic data set was imputed using quantile regression imputation for left censored data (QRILC) and principal component analysis was performed. A volcano plot was generated using “EnhancedVolcano”. ‘Pheatmap’ package was used to generate heatmap, clustering was performed using Euclidean distance for proteins and ward’s criterion (WardD2) for subjects. Gene set enrichment analysis (GSEA) was performed using the package “clusterprofiler” comparing against the Gene Ontogeny (GO) database with a P value of <0.5 deemed significant.

## Results

### Adult and neonatal mouse platelets have different proteomes

Our prior work found that adult and neonatal platelets had different mRNA expression, with adult platelets enriched in immune transcripts compared to neonatal platelets^21^. To determine whether these differences are reflected in the platelet proteomes, we isolated platelets from neonatal mice on post-natal day 1 (P1) and P3, as well as from adult mice (8-12 week old). To ensure a pure population of platelets after centrifugation-based isolation, we further depleted potential contaminating myeloid, lymphoid and erythroid cells with anti-CD45, CD220 and Ter119 beads. Mass spectrometry-based proteomics demonstrated that platelets from P1 and P3 neonatal mice had very similar proteomes (Supplementary Figure S1A-B). However, adult mice had a distinct proteome compared to the neonatal mice (Figure 1A-B). These findings were similar to studies that compared human adult and neonatal platelets^23^. Analysis of age dependent proteins further indicated that neonatal platelets were enriched for some growth factors, but deficient in many immune molecules compared with adult platelets (Figure 1B-C, representative proteins highlighted in C). In contrast P1 and P3 platelets were enriched in proteins related to ribosomes and mitochondria compared to adult (Figure 1D and Supplementary Figure S1D) perhaps indicative of more proliferative and metabolically active neonatal megakaryocytes and platelets. From these data, we concluded that P1 and P3 platelets have a similar proteome, but neonatal platelets diverge from that of adult platelets, not only in their mRNA expression, but also in their protein expression.

**Figure 1.**
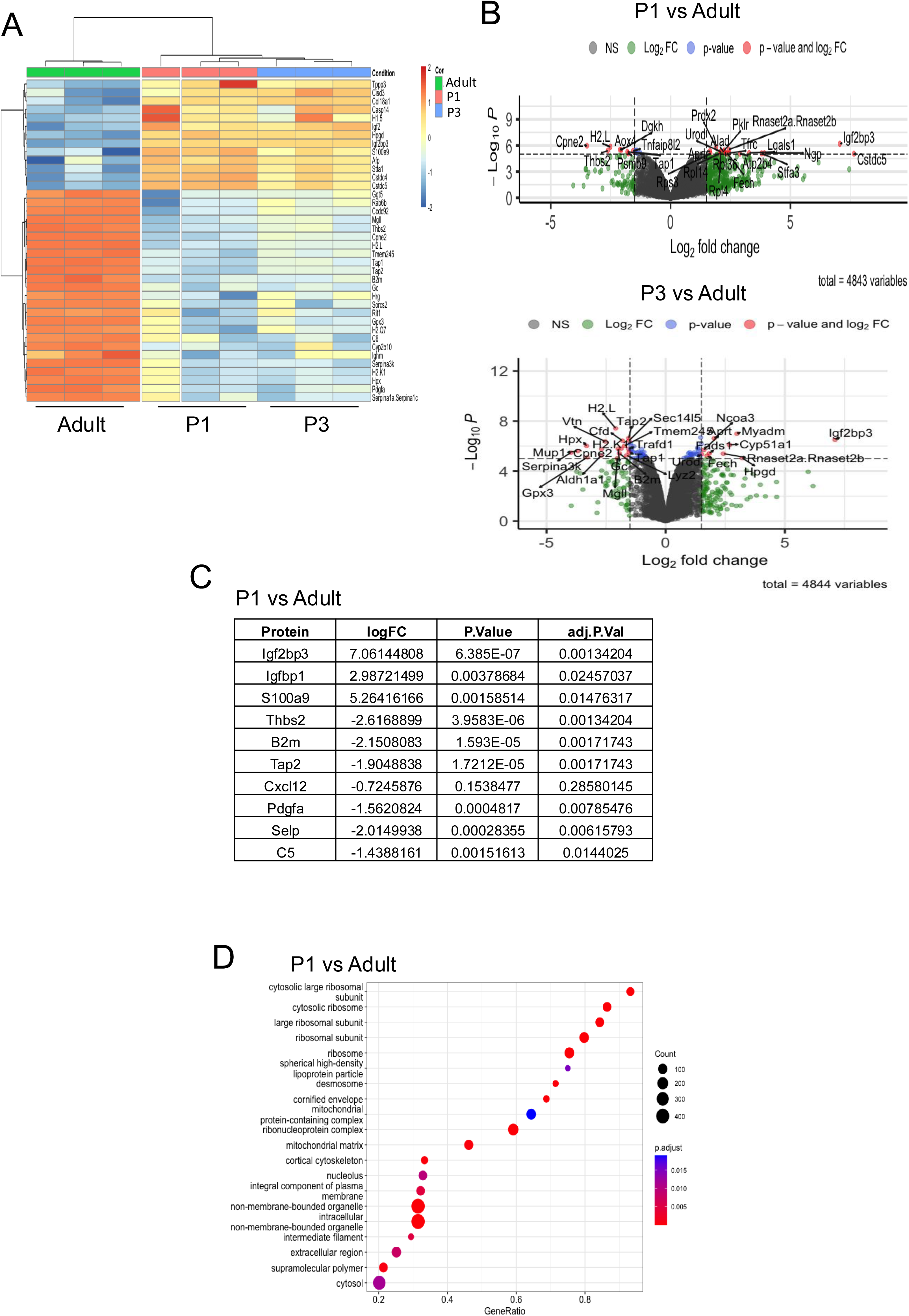
Neonatal and Adult Platelet Proteomes. P1 and P3 mouse platelet proteomes are similar, but neonatal platelet proteomes substantially differ from the adult platelet proteome. A) Heat map of adult, P1 and P3 platelet proteomes. B) Volcano plots of P1 or P3 vs adult platelet protein expression. C) Example of proteins changed in P1 vs Adult platelets. D) GO analysis of functional pathways more represented in P1 vs Adult platelets.

### Neonatal and adult platelets have different effects on monocyte differentiation

Our prior studies explored the acute and contact-dependent effects of neonatal platelets on monocytes^21^. To begin to explore whether neonatal and adult platelets have different effects on monocyte/macrophage differentiation, naïve adult bone marrow monocytes were treated for 48 hrs with releasates from adult or neonatal platelets that were prepared as we have prior published^20,24^ and monocyte mRNA was isolated for RNA-seq. Monocytes treated with neonatal or adult platelets had different gene expression profiles (Figure 2A). A deeper analysis of gene markers of monocyte and macrophage differentiation indicated that neonatal platelet releasates increased the expression of genes associated with myeloid derived suppressor cells (MDSC). This included a mix of both reparative (*Arg1, Vegfa, Il10*) and inflammatory (*Nos2, Il6*) genes, as well as genes more unique to MDSCs, including *Ptgs2, Ptges*, *Tnfrsf10b* (DR5), and *CD274* (PD-L1) (Figure 2B).

**Figure 2.**
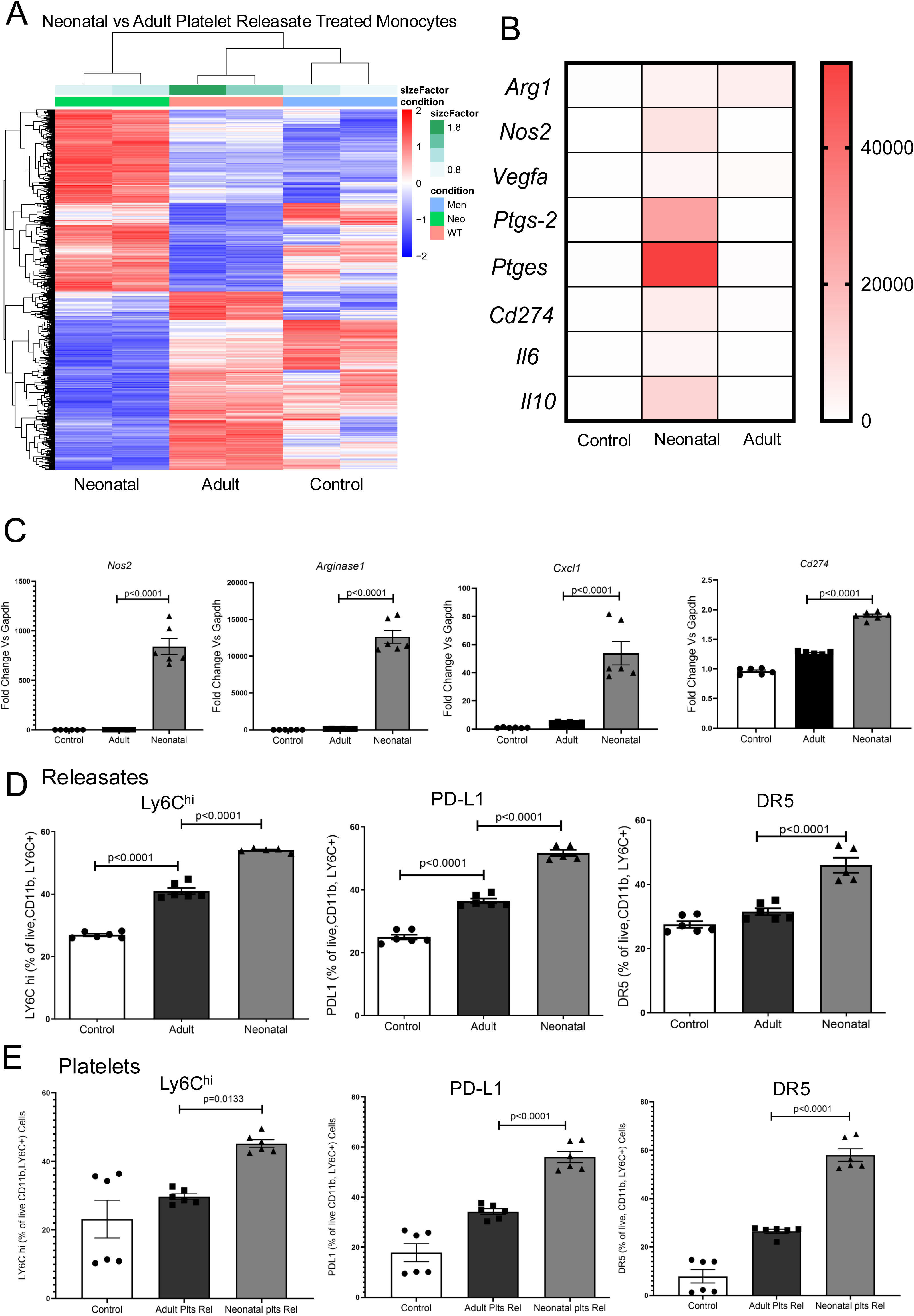
Neonatal platelet derived products induce a MSDC-like monocyte phenotype. Adult BM derived monocytes were incubated with control buffer, adult or P1-3 mouse platelet releasate for 48 hrs, and RNA-seq performed. A) Heat map of gene expression differences between monocytes treated with neonatal platelet releasates, adult platelet releasates, or control buffer. B) Relative expression of genes associated with a MDSC-like phenotype after treatments. C) Validation of MDSC associated genes using qRT-PCR after monocytes were treated with control, adult or neonatal platelet releasates for 48 hrs. D) Neonatal platelet releasates increased the expression of M-MDSC markers Ly6C, PD-L1 (CD274) and DR5 as determined by flow cytometry. E) Intact neonatal platelets increased the expression of M-MDSC markers Ly6C, PD-L1 (CD274) and DR5 as determined by flow cytometry. (Mean ± SEM, 1-way ANOVA with Bonferroni correction).

MDSC are a set of ‘immature’ myeloid cells that suppress T-cell responses and are present in limited numbers in healthy adults, but MDSCs may expand in some cancers where they are associated with immune therapy failure^28^. M-MDSC are present in cord blood but extremely rare in maternal blood and are typically identified by the markers CD11b^+^Ly6G^−^Ly6C^hi^PD-LI^hi^DR5^hi^^29,30^. MDSCs inhibit T-cell activation in a contact dependent manner through PD-L1 (CD274), a potent T-cell inhibitor, as well as via secreted molecules^31^. The same incubation conditions used for RNA-seq were used to analyze the expression of select M-MDSC-associated genes by qRT-PCR. *Nos2, Arg1, Cxcl1* and *Cd274* were all increased by neonatal platelet releasates (Figure 2C). M-MDSC specific markers were also quantified by flow cytometry. Neonatal platelet releasates increased the number of Ly6C^hi^PD-L1^hi^DR5^hi^ monocytes, consistent with a M-MDSC phenotype (Figure 2D). Furthermore, the incubation of intact neonatal platelets with monocytes also induced a M-MDSC phenotype (Figure 2E). We therefore concluded that the exposure of monocytes to neonatal platelets leads to an immune suppressive M-MDSC-like phenotype.

We next asked whether monocytes from neonatal mice expressed more MDSC associated molecules. Isolated adult and P3 monocytes had different patterns of gene expression, including neonatal monocytes having increased expression of MDSC associated genes (Supplementary Fig S2A-C). These data further indicate a propensity for increased MDSCs in neonates that may in part be platelet dependent.

### Neonatal Platelet Interactions with Monocytes Limit T-cell responses

Because M-MDSCs are potent inhibitors of T-cell activation, we next asked whether neonatal platelets indirectly limit T-cell activation by promoting monocytes to become MDSCs. Adult mouse bone marrow monocytes were isolated and cultured for 3d in control conditions or with adult or neonatal platelet releasates. On d3 T-cells were isolated from adult mouse spleens, T-cells labeled with CFSE dye, and added to the monocyte cultures in the presence of anti-CD3/28 antibody to activate T-cells in an antigen independent manner. 5d later CD4^+^ and CD8^+^ T-cell proliferation was measured by CFSE dye dilution (example gating in Supplementary Figure S3), and T-cell activation measured by CD25 and CD69 expression (Figure 3A). The prior incubation of monocytes with neonatal platelet releasate limited CD4^+^ (Fig 3B) and CD8^+^ (Fig 3C) T-cell proliferation and activation. MDSCs can promote an expansion of CD4^+^ T-cell differentiation to T regulatory cells (Tregs) that have an immune limiting role^32^. However, we found that the prior incubation of monocytes with neonatal platelet releasate limited Tregs as determined by intracellular Foxp3 (Supplementary Figure 3B), likely due to the decreased T-cell activation. From these data, we concluded that neonatal platelet releasates induced M-MDSCs differentiation that limited T-cell proliferation and activation, perhaps providing a more immune tolerant environment in the post-natal period.

**Figure 3:**
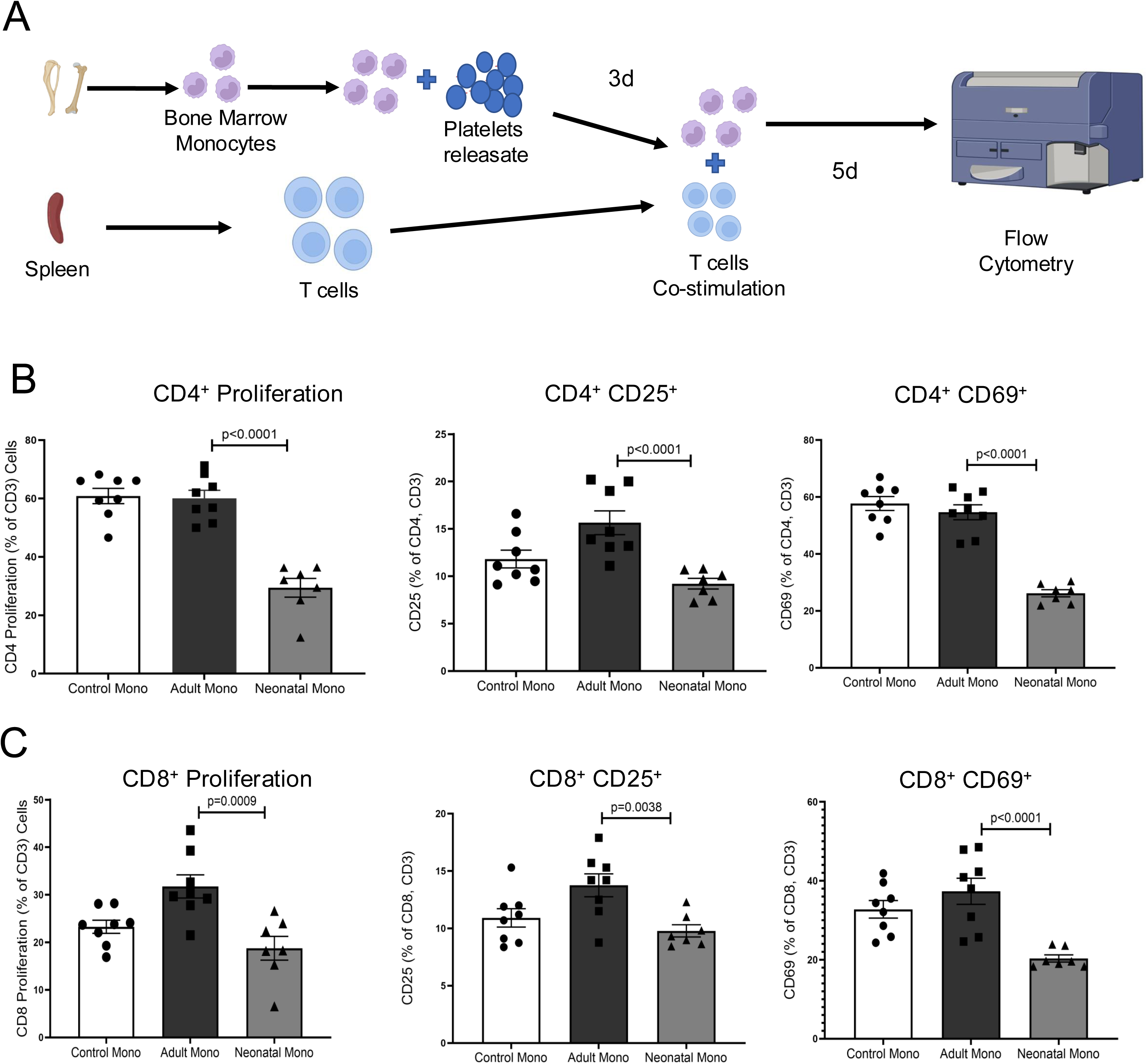
**Neonatal platelet induced MDSCs limit T-cell activation *in vitro.*** Adult BM derived monocytes were incubated with adult or P1-3 platelet releasate for 3d prior to the addition of adult spleen isolated T-cells. T-cells were activated with anti-CD3/CD28 antibody and 5d later flow cytometry for T-cell proliferation (CFSE dilution) and activation (CD25 and CD69) were determined. A) Study design schematic. B-C) Monocytes prior incubated with neonatal, but not adult platelet releasates reduced B) CD4^+^ and C) CD8^+^ T-cell proliferation (CFSE dye dilution) and activation (CD25 and CD69 expression). (Mean ± SEM, 1-way ANOVA with Bonferroni correction).

To determine whether adult monocytes exposed to neonatal platelet releastes limit T-cell responses *in vivo*, we utilized an antigen-induced lung inflammation model. Adult monocytes were incubated with adult or neonatal platelet releasates for 3d and monocytes then transfused into adult mice (PBS only as control) on experimental d0 and again on d6. Mice were challenged intraperitoneal (i.p.) with ovalbumin (OVA) on d1 and d7 to prime T-cell responses. On d14-16 mice were OVA challenged oropharyngeal (o.p.) to induce an antigen-specific lung T-cell response (PBS as no stimulation control, Figure 4A). There were increased total immune cells (CD45^+^ cells) in mice exposed to only OVA or OVA and adult platelet co-incubated monocytes (Supplementary Figure S4A). However, CD45^+^ cell infiltrates in response to OVA were limited by the prior administration of monocytes co-incubated with neonatal platelet releasates (Supplementary Figure S4A). These data were reflected in hematoxylin and eosin stained lungs sections (Supplementary Figure S4B). The percentage of CD3^+^CD45^+^ cells was similar in all groups compared to control mice, except the neonatal platelet/monocyte treated group, which was slightly increased (Supplementary Figure S4C). However, T-cell activation was limited in the neonatal platelet/monocyte group as demonstrated by CD4^+^ (Figure 4B) and CD8^+^ (Figure 4C) T-cells from the lung having reduced CD25 and CD69 expression compared to the other OVA treated groups. Because MDSCs limit T-cell activation, there are fewer T-helper type 1 (Th1, T-bet^+^ cells) and Th2 (GATA-3^+^ cells) T-cells in the lungs of mice given neonatal platelet treated monocytes (immunohistochemistry Figure 4D quantified in Supplementary Figure S4D-E), despite flow cytometry showing similar numbers of total T-cells in all groups of mice. Taken together, these data suggest that neonatal platelets induce monocytes to become MDSCs that limit T-cell activation both *in vitro* and *in vivo*.

**Figure 4:**
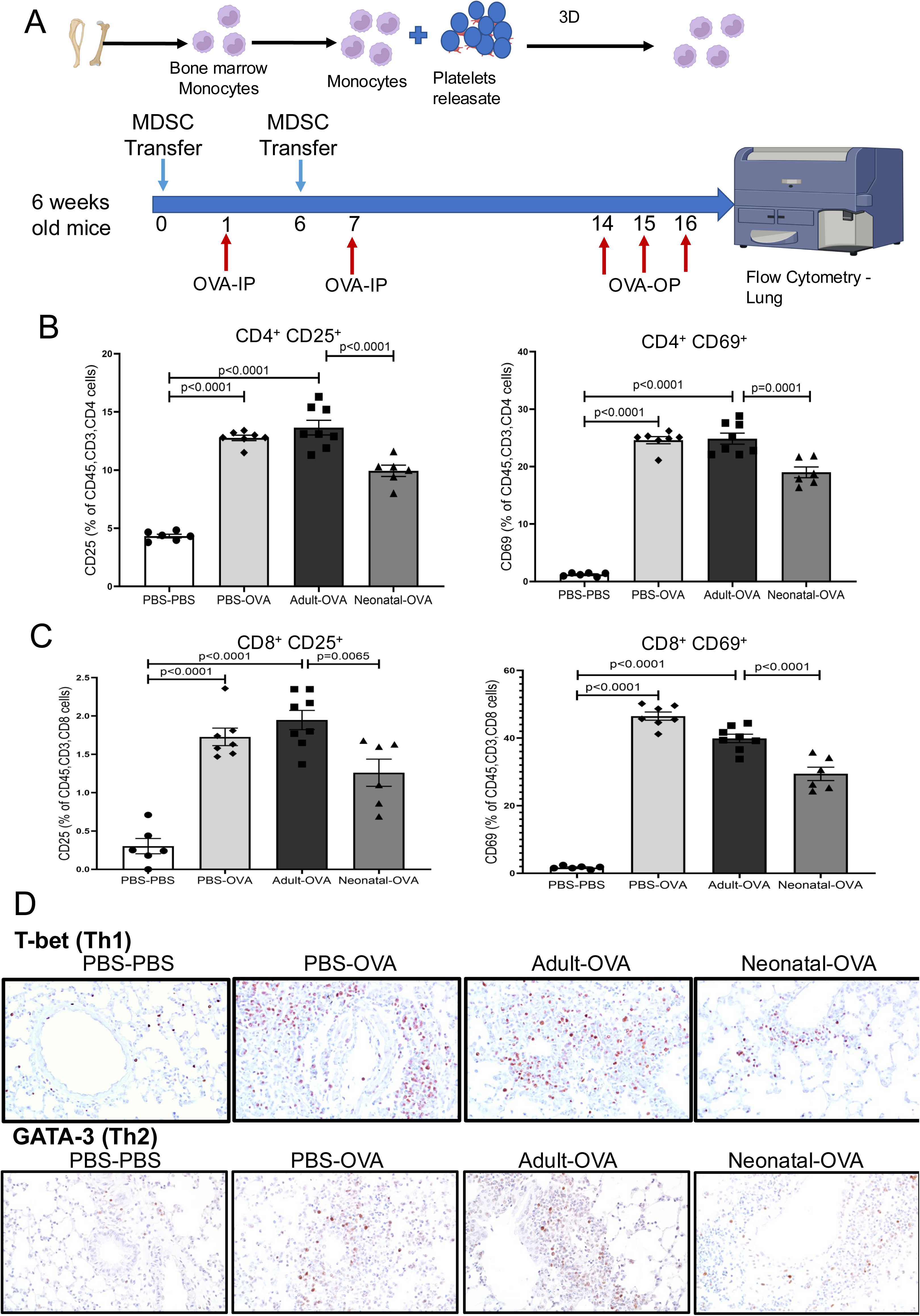
**Transfer of neonatal platelet derived M-MDSCs limits T-cell responses *in vivo.*** Adult BM derived monocytes were incubated with PBS, adult or P1-3 mouse platelet releasate for 3d prior to i.v. injection into adult mice. 1 and 7d after the first monocyte injection mice were given OVA i.p. to prime a T cell response and on d14, 15, 16, OVA o.p. to drive a T-cell response in the lung. 1d after last OVA treatment mice were euthanized and lung cell infiltrates quantified by flow cytometry. A) Experimental design schematic. B-C) Mice given monocytes after incubation with neonatal platelet releasates had less B) CD4^+^ and C) CD8^+^ T-cell activation. D) Immunostains for markers of Th1 (T-bet) and Th2 (GATA-3) T-helper cells. (Mean ± SEM, 1-way ANOVA with Bonferroni correction).

### Neonatal platelet induced MDSC differentiation is at least in part PGE_2_ dependent

We next sought to determine the mediators of neonatal platelet-dependent MDSC differentiation. We first tested IGFBP2 which is increased in neonatal platelet proteome and has been implicated in promoting immunosuppressive macrophages^33^. IGFBP2 had no effect on MDSC differentiation (Supplementary Figure S5A). Prostaglandin E_2_ (PGE_2_) is a described mediator of MDSC differentiation^34,35^ and peptides related to enzymes that synthesize PGE_2_ (Ptges2 and Ptges3) had increased abundance in the proteomics data (Supplementary Figure S5B). Immunoblot of adult and neonatal platelets confirmed that neonatal expressed more of a major PGE_2_ synthesis enzyme PGES-1 compared to adult platelets (Figure 5A). This was further confirmed by immunofluorescent imaging of platelets (DAPI negative cells as platelet confirmation, Supplementary Figure S5C). We therefore performed PGE_2_ ELISA on releasates from the same number of activated adult and P3 platelets and found that neonatal platelets produced significantly more PGE_2_ compared with adult platelets (Figure 5B). The cell culture supernatant from monocytes incubated with neonatal platelets also had more PGE_2_ compared to those cultured with adult platelets (Figure 5B). To provide a mechanistic link from neonatal platelet PGE_2_ to MDSC differentiation we incubated neonatal (Figure 5D) or adult monocytes (Supplementary Figure S5D) with PGE_2_. Recombinant PGE_2_ increased the expression of PD-L1 and DR5 in both neonatal and adult monocytes in a dose dependent manner very similar to neonatal platelet releasates (Figure 5D and Supplementary Figure S5D). PGE_2_ signals through EP2 and EP4^36^. To more directly demonstrate PGE_2_ dependent signaling, we blocked either EP2 or EP4 prior to addition of neonatal platelet releasates. Blocking EP2 had some effect on neonatal platelet driven MDSCs (Supplementary Figure S5E), while EP4 blockade lead to a significant decrease in Ly6C, PD-L1, and DR5 expression (Figure 5E, dose response in Supplementary S5F). Furthermore, pre-treatment of neonatal platelets with the cyclooxygenase inhibitor indomethacine prior to activation greatly attenuated neonatal platelet induced MDSC differentiation (Figure 5F). We therefore conclude that PGE_2_ from neonatal platelets in part mediates the differentiation of monocytes to a MDSC phenotype.

**Figure 5.**
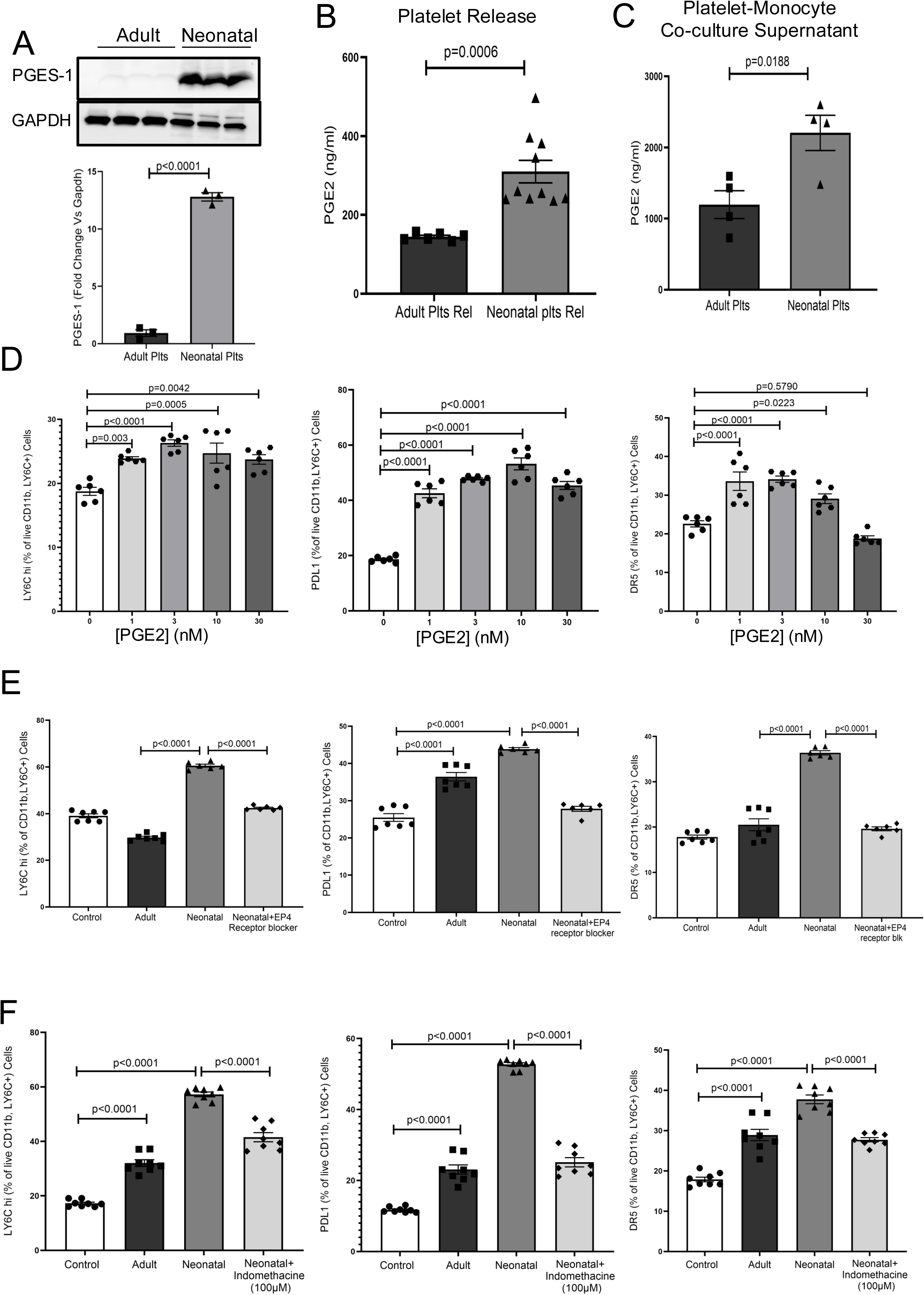
Neonatal platelet derived PGE_2_ is a mediator of MDSC differentiation. A) Neonatal platelets express more PGES-1 compared to adult platelets. Immunoblot and quantification. B) Activated neonatal platelets release more PGE_2_ compared to adult platelets. C) Cell culture supernatants from monocyte-neonatal platelet co-culture had more PGE_2_. D) Exogenous PGE_2_ increased neonatal monocyte MDSC differentiation. E) EP4 receptor blockade limited neonatal platelet induced M-MDSC differentiation. F) Incubation of neonatal platelets with the cyclooxygenase inhibitor indomethacin reduced neonatal platelet mediated MDSC differentiation (Mean ± SEM, 1-way ANOVA with Bonferroni correction).

## Discussion

Adult platelets are more enriched in immune molecules than are neonatal platelets^21,22^, which may impact neonate responses to platelet transfusions. Our past study using a thrombocytopenic neonatal mouse model showed that adult platelet transfusions induced monocyte trafficking while neonatal platelets did not^21^. Our new data extend our knowledge of the immune regulatory roles for neonatal platelets to include limiting adaptive immune responses in the neonate, by facilitating monocyte differentiation to a MDSC phenotype. This conceptually makes sense, as the fetus must avoid rejection by the maternal immune system, and the post-natal period is a time of significant microflora colonization as well as tissue remodeling and continued development. It is therefore important to maintain a relatively quiescent adaptive response when the organism is exposed to many new antigens.

Our data indicates that PGE_2_ is a mediator of neonatal platelet driven M-MDSC differentiation. PGE_2_ can be produced by most cells of the body and impacts multiple types of immune cells as well as endothelial cells, but the physiologic consequences depend on the cell type, PGE_2_ concentration, and the tissue environment. PGE_2_ acts on EP receptors limiting macrophage phagocytosis and PGE_2_ supports the induction of mature dendritic cells that are capable of homing to lymph nodes to prime naïve T-cells^37,38^. PGE_2_ at high concentrations has direct effects on T-cells by non-specifically suppressing T-cell activation and proliferation. However, at low concentrations PGE_2_ shifts differentiation of activated CD4^+^ T-cells from Th1 cells to Th2 and Th17 cells^39,40^. It is well established that in mammalian neonates a Th2 type response predominates to facilitate the establishment of the microbiome in postnatal life and prevent preterm birth or immunologic rejection by the maternal immune system^1^. Prolonged maintenance of Th2 polarity has been implicated in the development of atopy and allergy^41,42^. PGE_2_ therefore has complex roles in immune differentiation and responses, which are even more multi-dimensional in the neonate. PGE_2_ is ussed clinically to maintain patency of the ductus arteriosus in neonates with “duct dependent” cyanotic congenital cardiac defects. Current medical treatments for an inappropriately patent ductus arteriosus (PDA) target either cyclooxygenases or perioxidases to reduce PGE_2_ and facilitate duct closure. Increased neonatal platelet generation of PGE_2_ is likely to have biologic functions beyond these findings.

Because many immune molecules and growth factors differ between neonatal and adult platelets, both increased and decreased, it is likely that PGE_2_ is one of many immune mediators in neonatal platelets that may affect all phases of immune responses. Our studies utilized mouse models to gain a mechanistic understanding of platelet directed developmental immunology that we hope can be translated to future clinical studies. Studies using human platelets have also shown that human neonatal platelets phenocopy mouse data and are relatively deficient in immune molecules compared to adult^23^, but further study is needed to extend these findings in more clinical directions, particularly in the context of transfusions. Of note, these studies only address the indirect effects of platelets on T-cell activation, not more direct platelet effects or how platelets may influence T-cell differentiation. Those questions can now build from these data to develop a more comprehensive understanding of neonatal platelet immune regulation and its impact on childhood health and disease.

Whether complications associated with platelet transfusions to neonates, particularly those more long-term, may be in part driven by increased T-cell activation and differentiation in association with a reduction in MDSCs requires more clinical and animal model-based study. There are clinical reports of reduced PGE_2_ being associated with asthma^43,44^, perhaps through direct T-cell effects or indirectly via reductions in MDSCs. Delivery of PGE_2_ concomitant with platelet transfusions to neonates may help to resolve some transfusion complications, but the dose and other physiologic implications would need extensive study to fully understand the best approach especially given the well-recognized complexity of PGE_2_ targets and signaling.

## Acknowledgements

CNM received funding from NIH/NHLBI, HL160610, HL153409

## Author Contributions

PM developed ideas, performed experiments, analyzed data and contributed to writing, DOR performed experiments, analyzed data and contributed to writing, ZTH performed experiments, analyzed data, KEM analyzed data, ACL, CL, MM, ETT, and SKT performed experiments, JP analyzed data and contributed to writing, CNM developed ideas, performed experiments, analyzed data and contributed to writing

## Conflict of Interest

All authors declare no competing financial interests.

## Data Sharing

For data requests, please contact. Proteomics data will be deposited in a public database (*PRIDE*) upon manuscript acceptance or made directly available with reasonable request.

## Key points

- Neonatal platelets differentiate monocytes to myeloid derived suppressor cells
- PGE_2_ is a mediator of neonatal platelet driven MDSC differentiation

## Notes

### Competing Interest Statement

The authors have declared no competing interest.

