## Supplemental data for "Neonatal Platelets Differentiate Monocytes to a Myeloid Derived Suppressor Cell Phenotype"

### Supplementary Figure S1

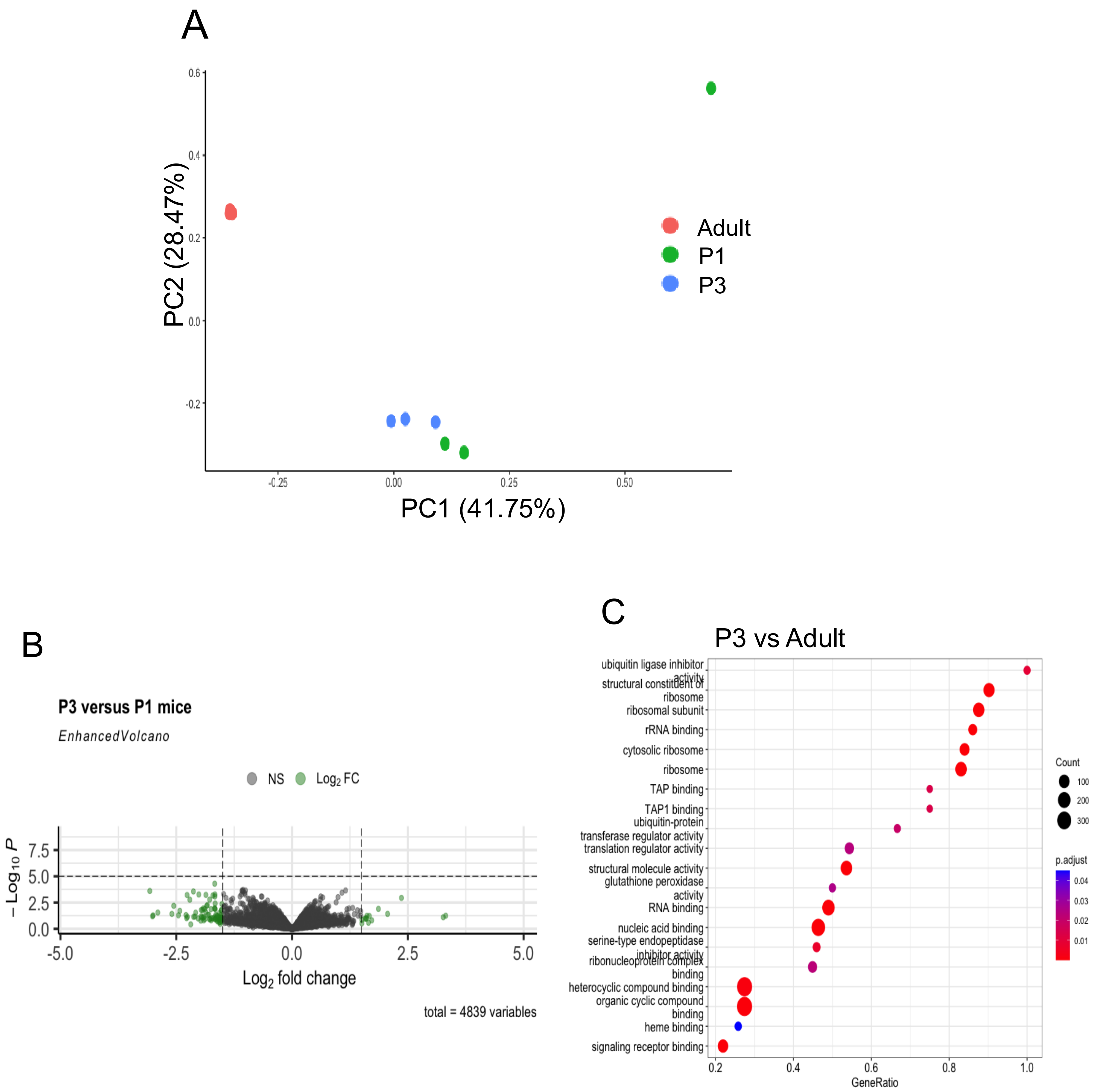


Supplementary Data S1. Platelet proteomics. A-B) The platelet proteome of P3 and P1 mice are very similar, but diverge from adult platelets A) PCA plot, B) P3 vs P1 volcano plot. C) GO pathway analysis of protein expression in P3 vs Adult platelets.

### Supplementary Figure S2

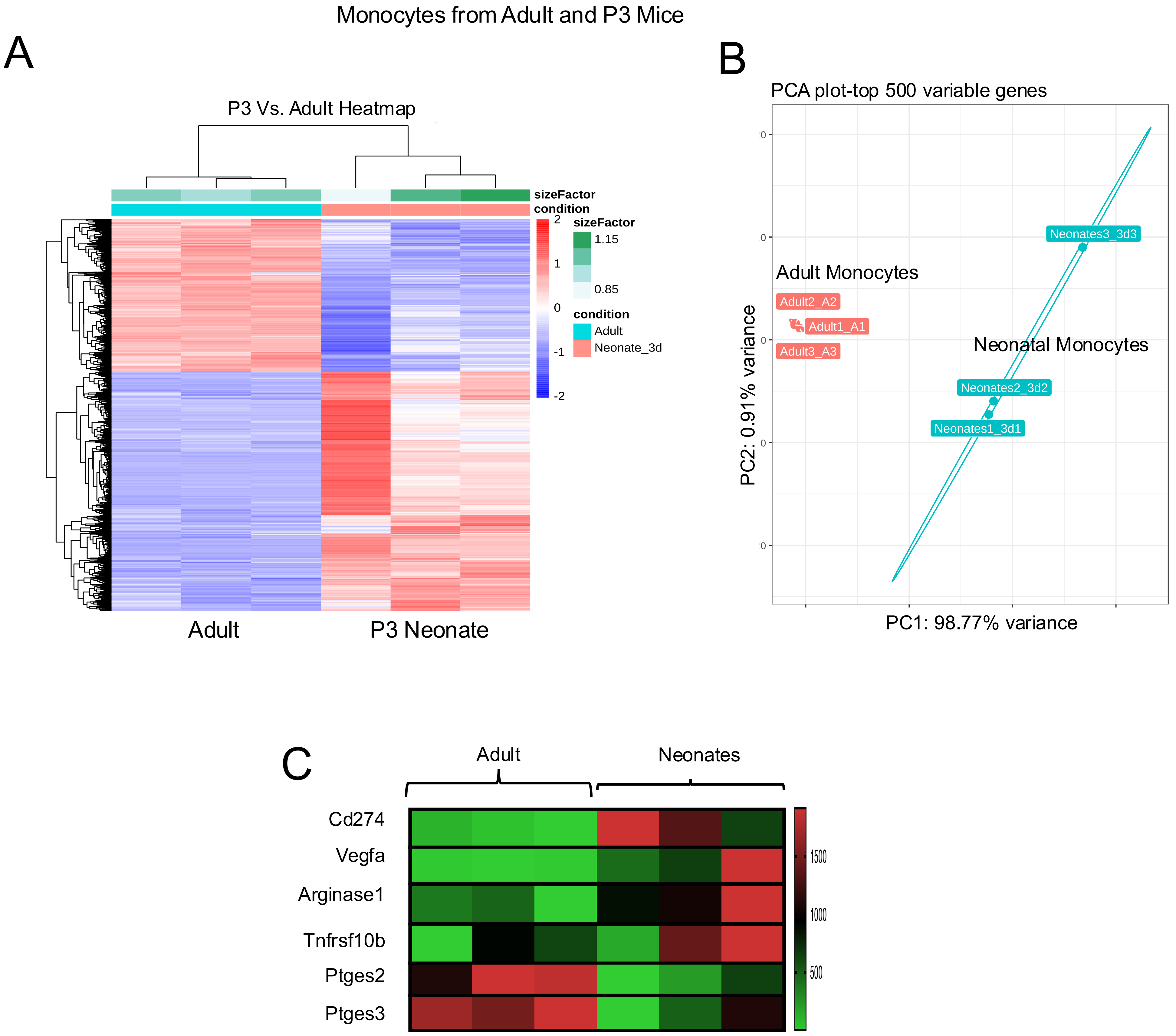


Supplementary Data S2. RNA-seq on monocytes from the BM of adult and P3 mice. P3 mouse monocytes have gene expression similar to M-MDSCs. A) Heat map and B) PCA plot demonstrate distinct gene expression patterns between P1 and adult monocytes. C) Relative expression of genes indicative of a MDSC phenotype.

### Supplementary Figure S3

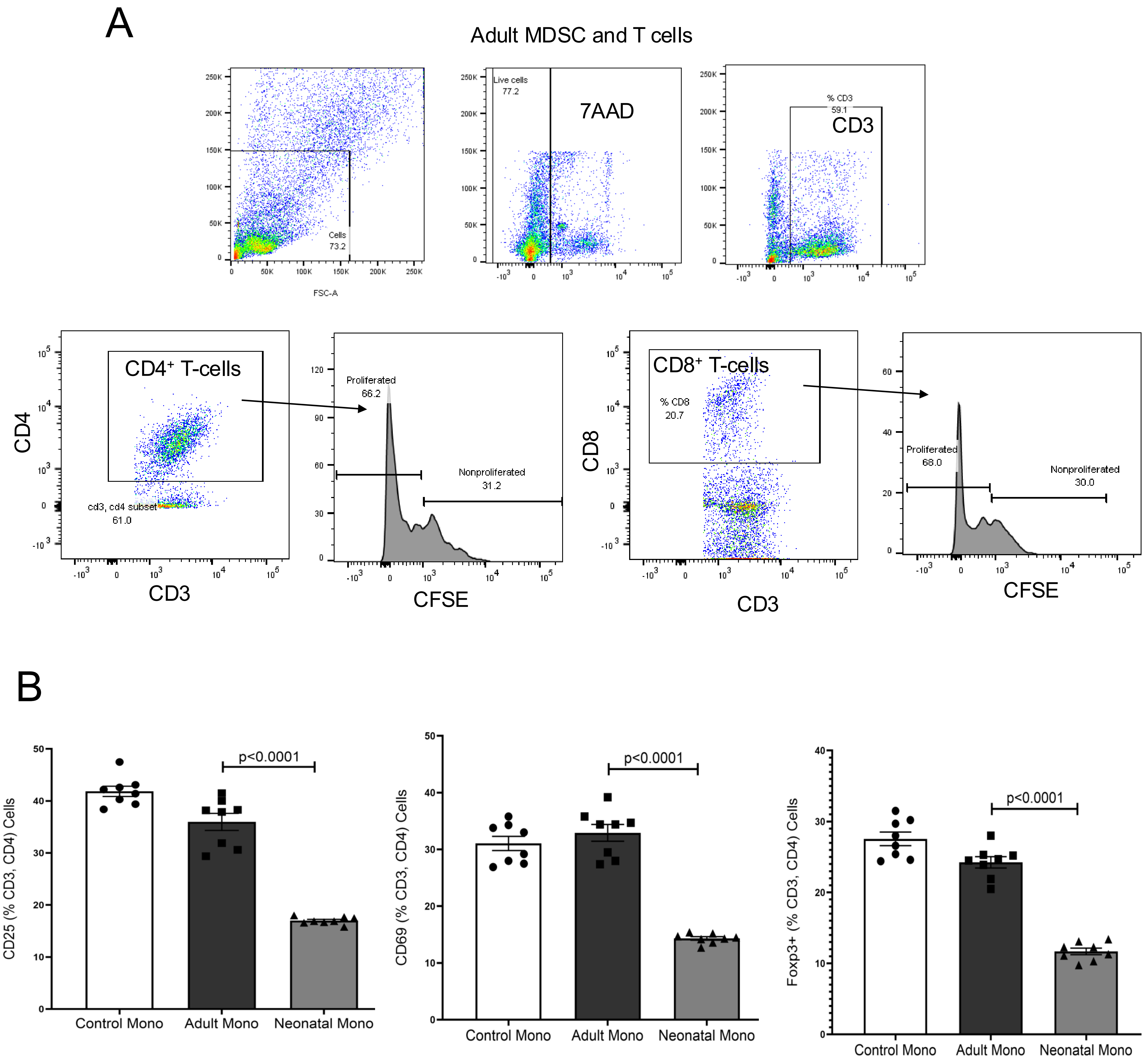


Supplementary Data S3. A) Representative flow cytometry gating to assess T-cell responses *in vitro.* Proliferation determined by CFSE dye dilution. B) Monocytes prior incubated with neonatal platelet releasates limited CD4⁺ T-cell activation as measured by CD25 and CD69. Foxp3 expression, indicative of Tregs, was limited in T-cells activated in the presence of monocytes prior incubated with neonatal platelet releasates, as determined by intracellular flow cytometry.

### Supplementary Figure S4

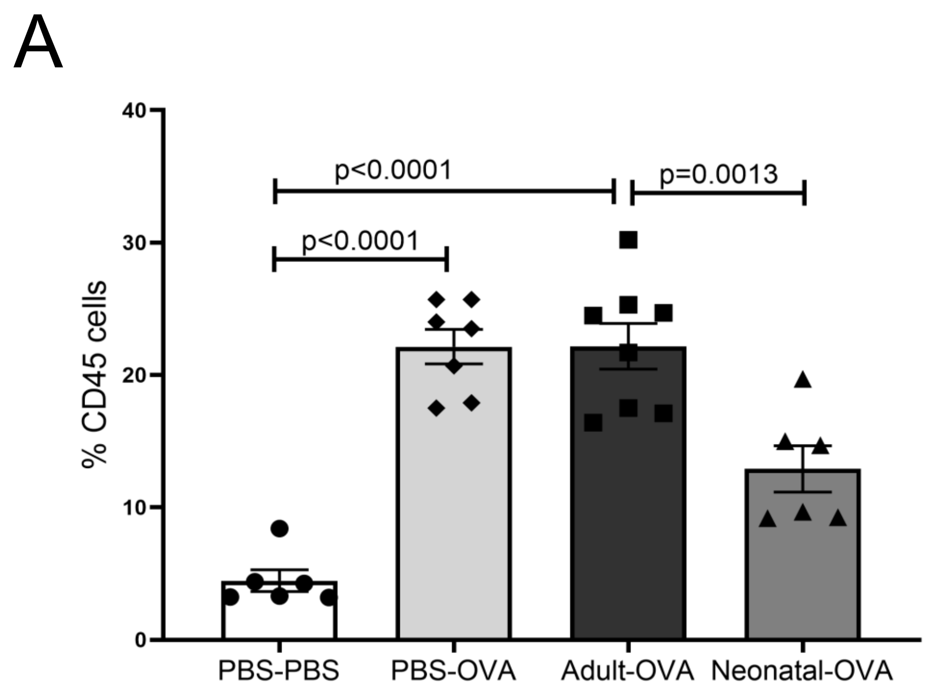


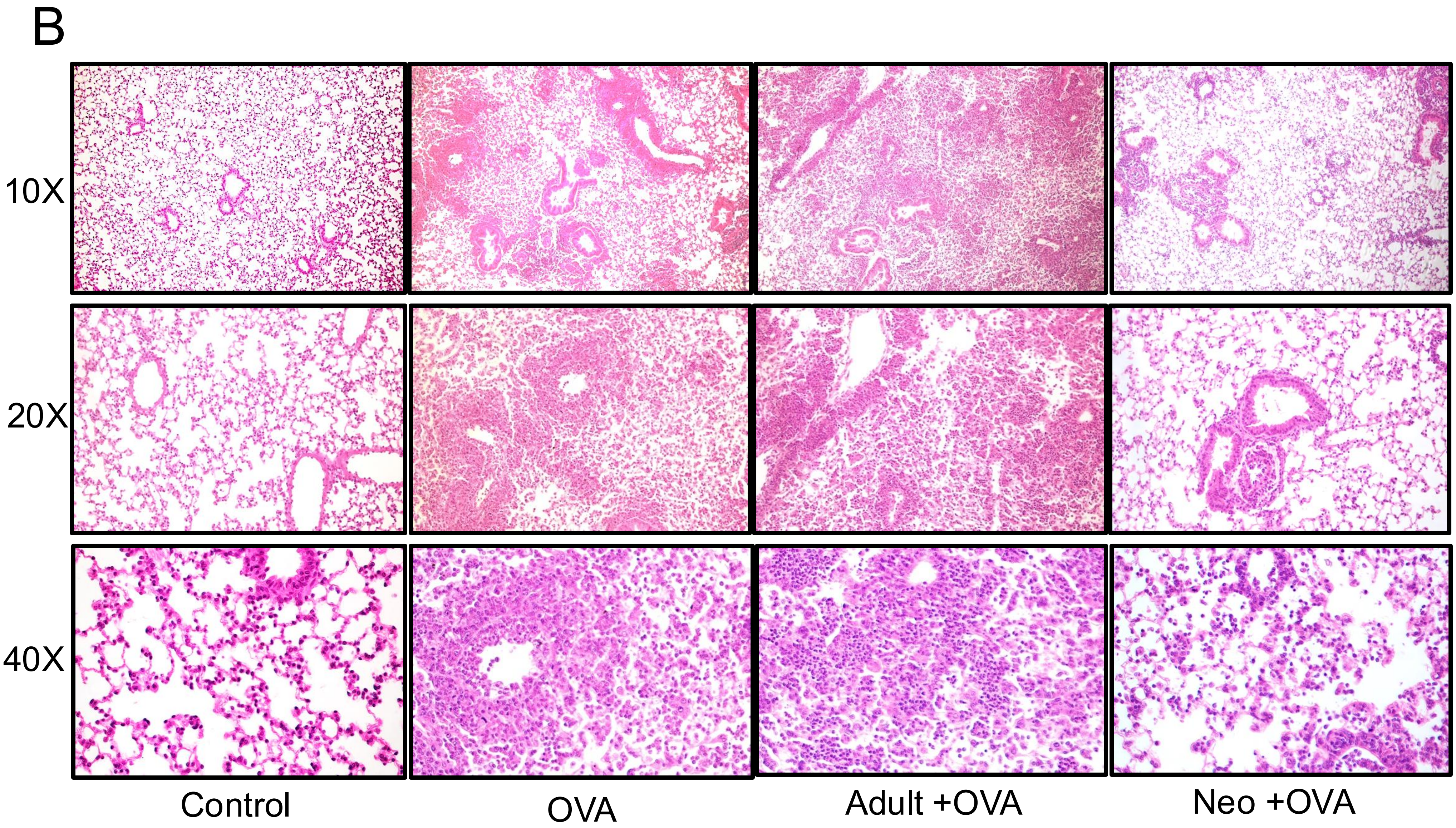


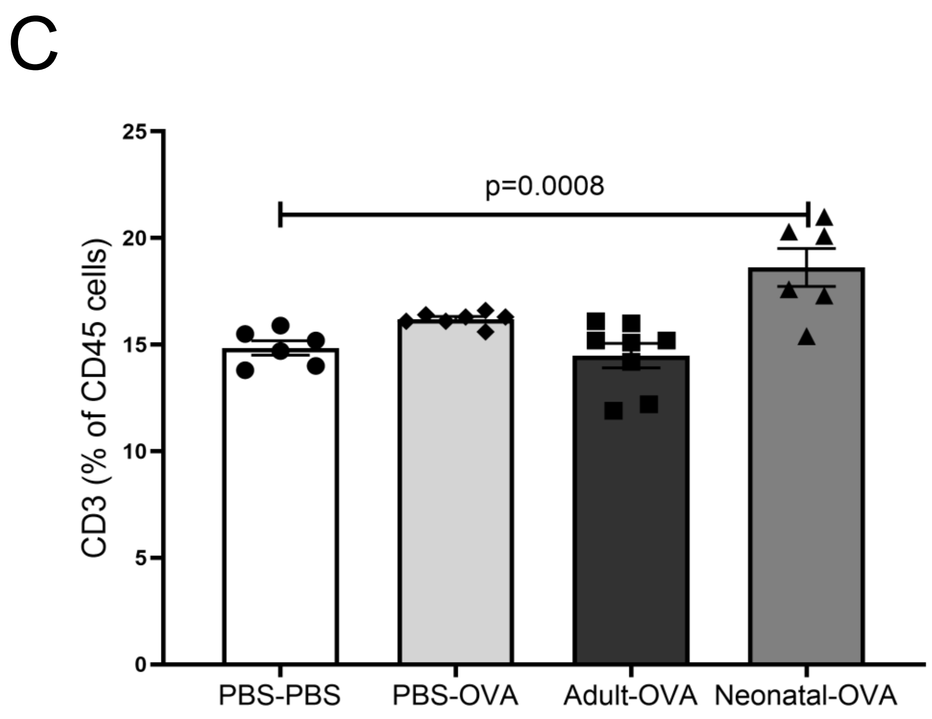

D

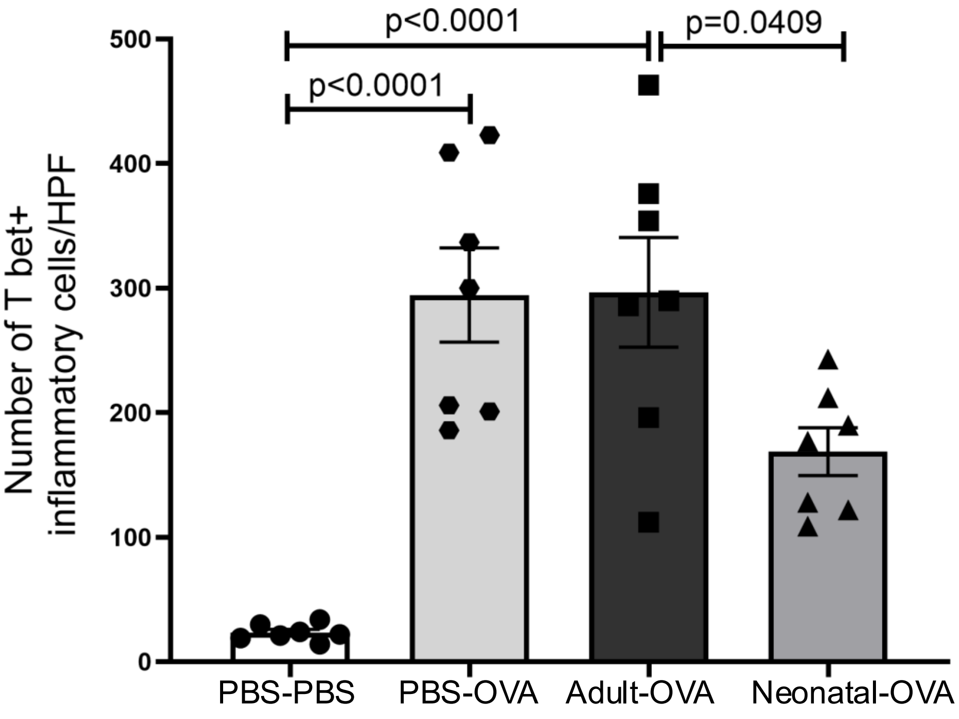


E

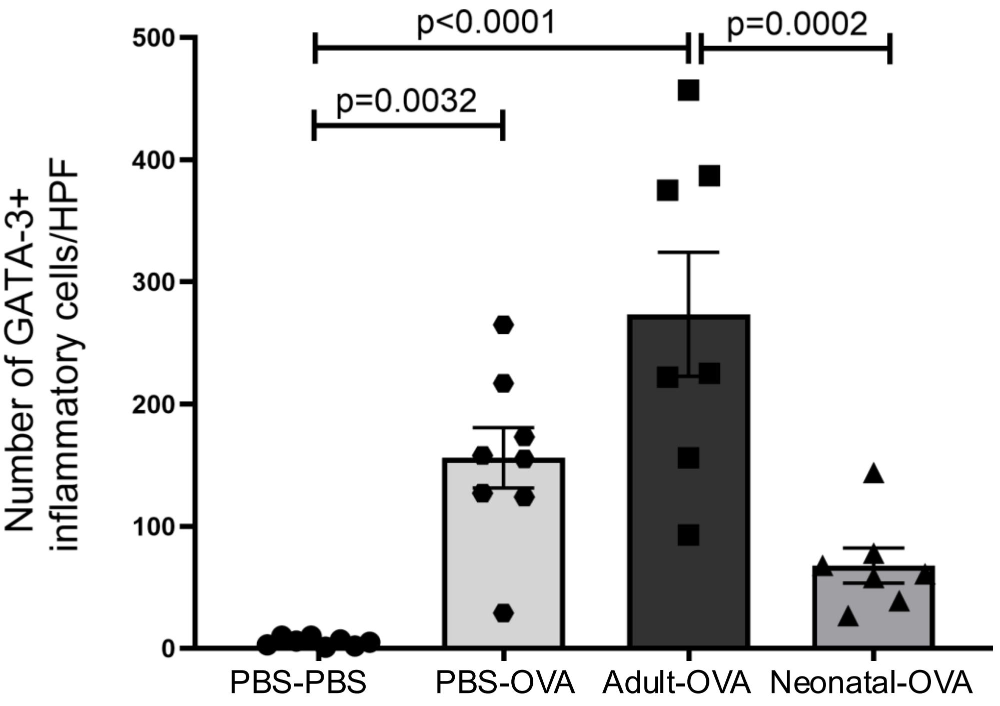


Supplementary Data S4. Neonatal platelet induced MDSCs limited antigen dependent T-cell responses in the lung. A) $CD45^+$ lung cells. B) Examples of lung histology. C) $CD3^+$ lung cells. D-E) Immunohistochemistry quantification of D) T-$bet^+$ (Th1) and E) GATA-$3^+$ (Th2) cells in the lungs.

### Supplementary Figure S5

A

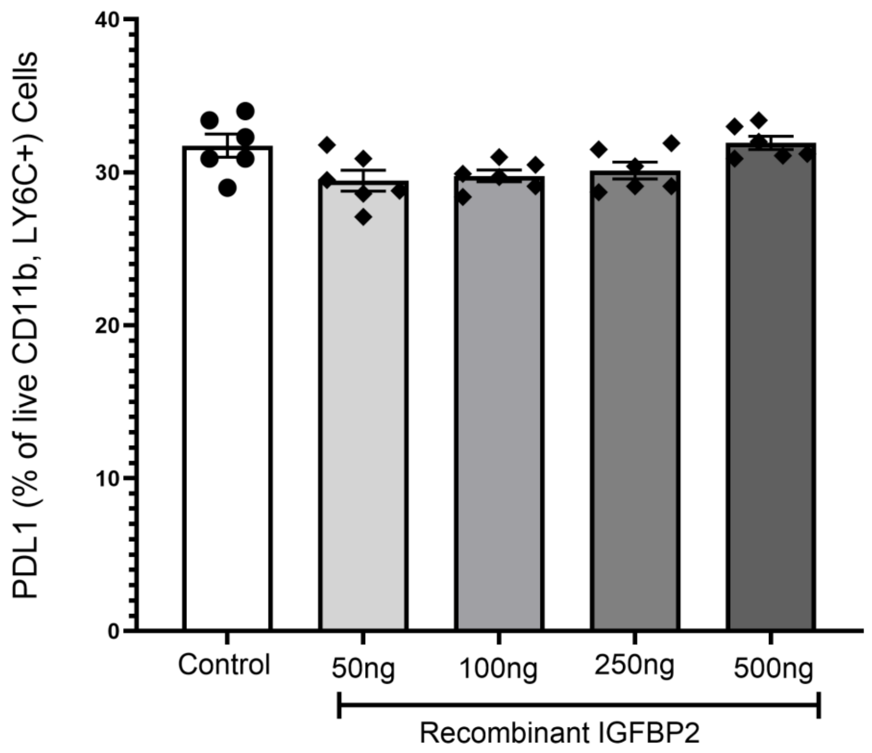


B

Median peptide abundance

| Protein | Adult | P3 |
|---|---|---|
| Ptges2 | 7.30E+04 | 8.57E+04 |
| Ptges3 | 1.25E+06 | 1.37E+06 |

C

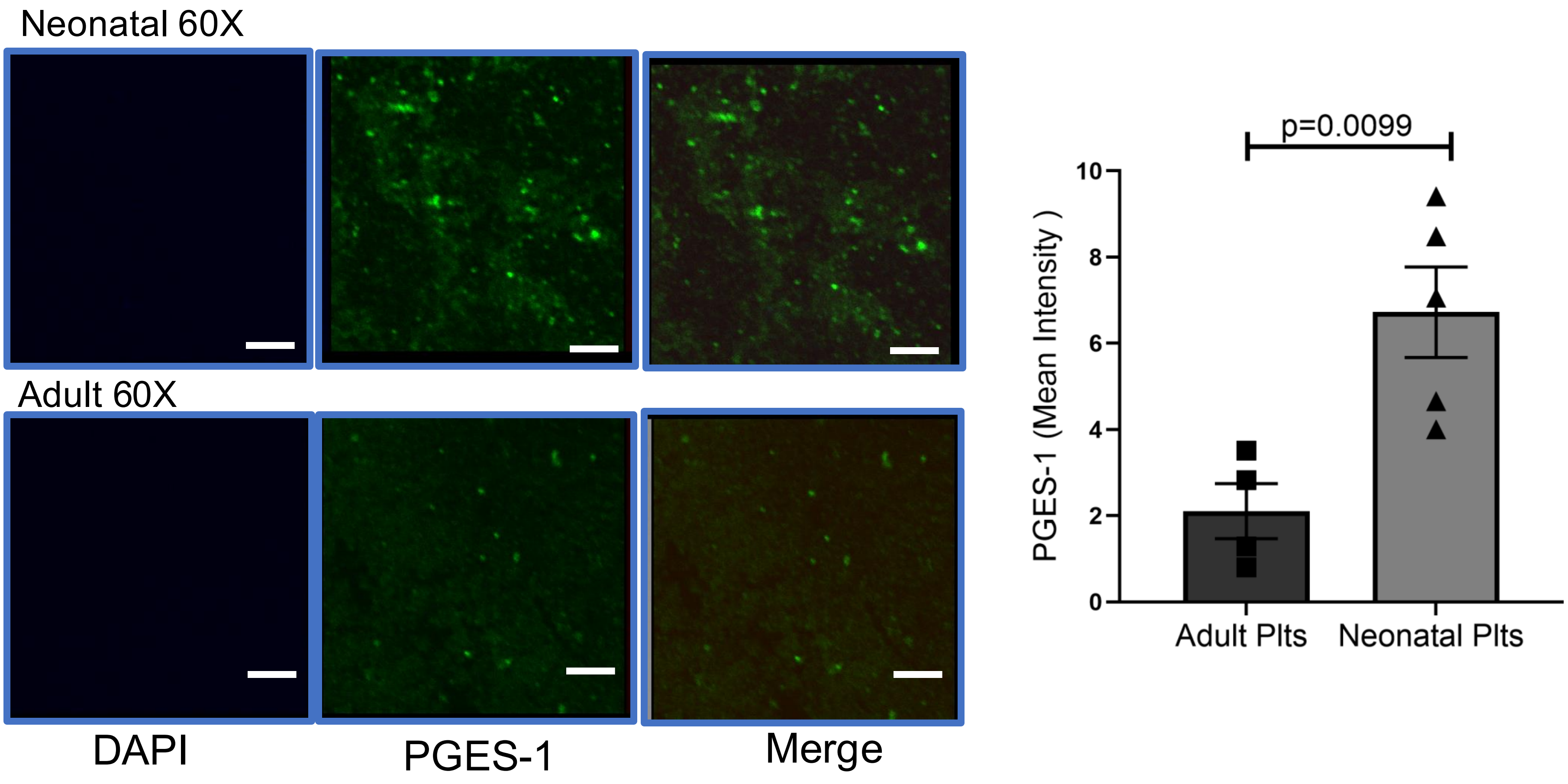


D

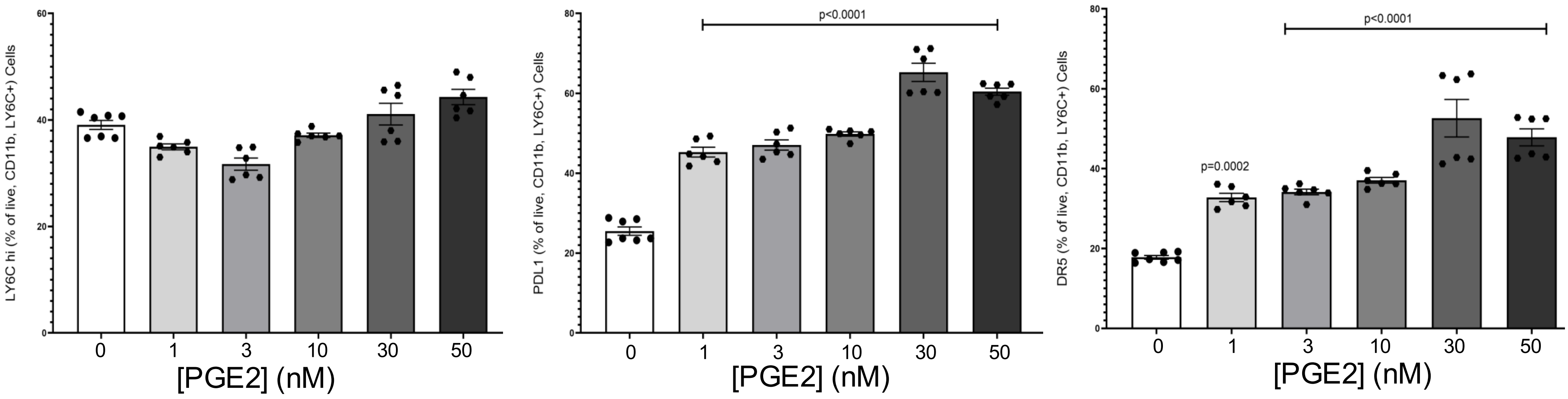

E

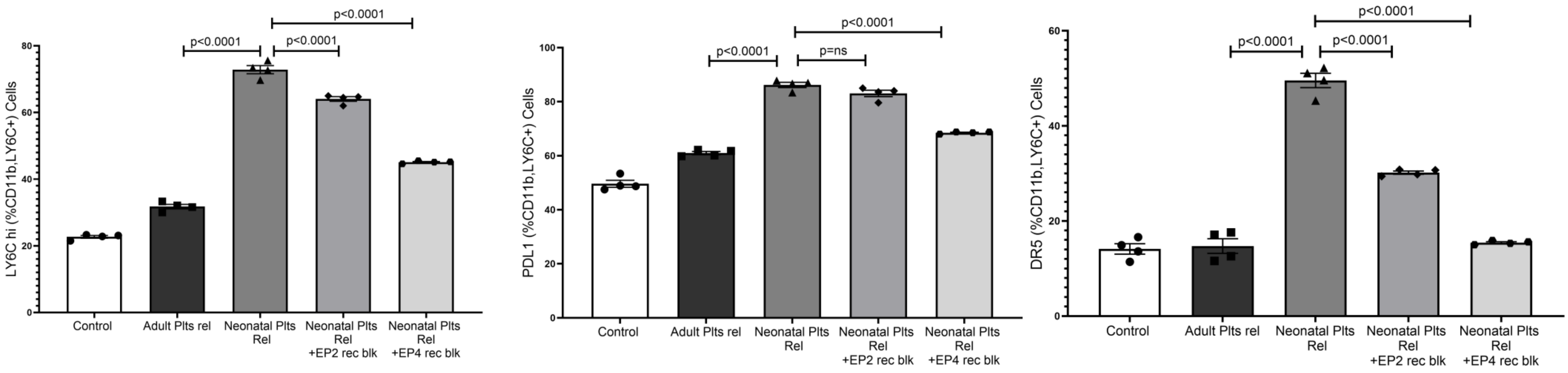


F

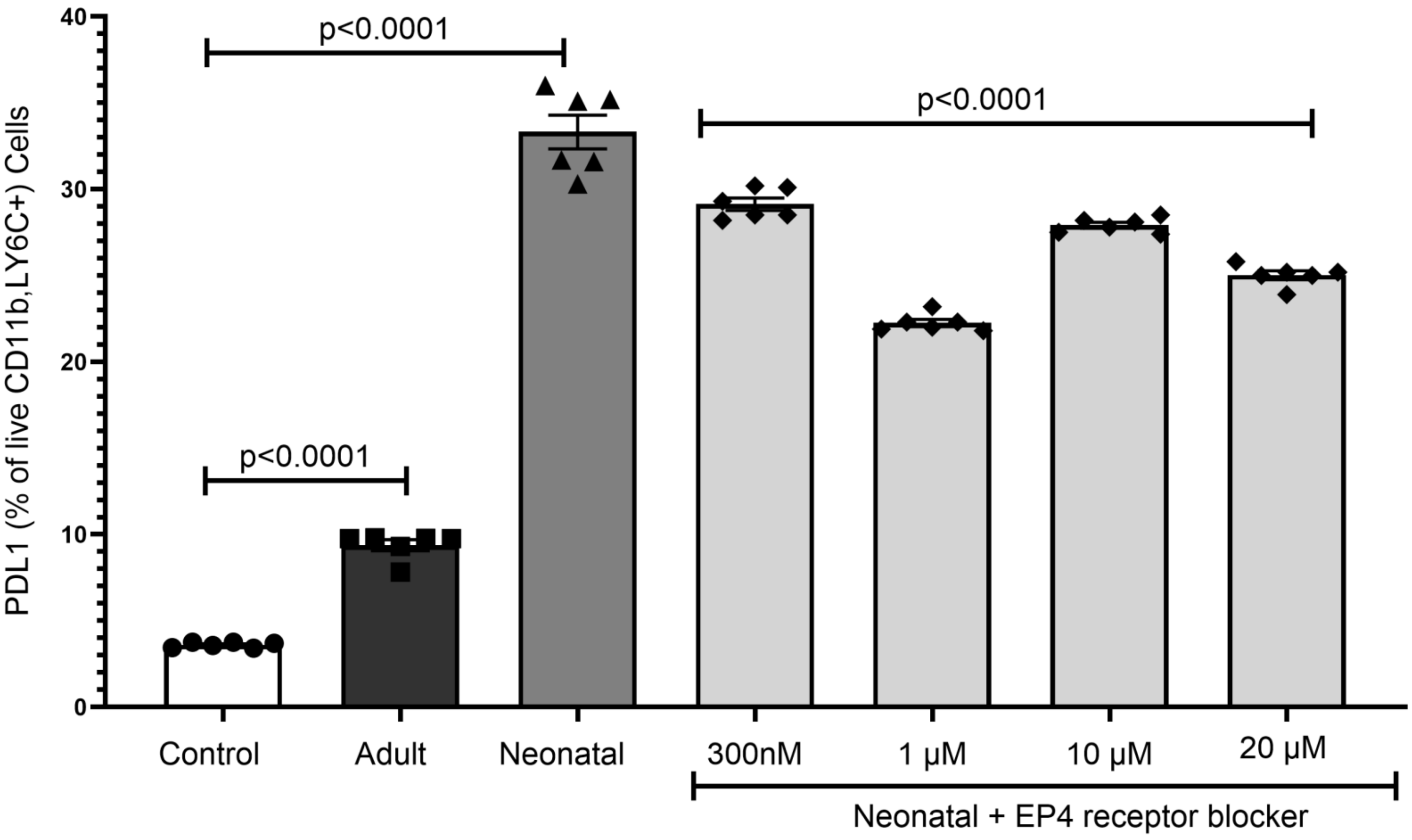


Supplementary Data S5. A) Recombinant IGFBP2 does not increase monocyte PD-L1 expression. B) Prostaglandin synthesis related proteins in proteomics data. C) Fluorescent imaging of PGES-1 in neonatal and adult platelets. D) Exogenous $PGE_2$ added to adult monocytes phenocopies neonatal platelet releasate effects on MDSC differentiation. E) Blocking monocyte EP4 has a greater effect on M-MDSC differentiation compared to EP2 blocking. F) Dose response with EP4 blocker and neonatal platelet releasate addition (Mean ± SEM, 1-way ANOVA with Bonferroni correction).
